# Expression of a recombinant, 4’-Phosphopantetheinylated, active *M. tuberculosis* Fatty acid Synthase I in *E. coli*

**DOI:** 10.1101/400622

**Authors:** Szilvia Baron, Yoav Peleg, Jacob Grunwald, David Morgenstern, Nadav Elad, Moshe Peretz, Shira Albeck, Yishai Levin, John T. Welch, Kim A. Deweerd, Alon Schwartz, Yigal Burstein, Ron Diskin, Zippora Shakked, Oren Zimhony

## Abstract

Fatty acid synthase 1 (FAS I) from *Mycobacterium. tuberculosis* (*Mtb*) is an essential protein and a promising drug target. FAS I is a multi-functional, multi-domain protein that is organized as a large (1.9 MDa) homohexameric complex. Acyl intermediates produced during fatty acid elongation are attached covalently to an acyl carrier protein (ACP) domain. This domain is activated by the transfer of a 4’-Phosphopantetheine (4’-PP, also termed P-pant) group from CoA to ACP catalyzed by a 4’-PP transferase, termed acyl carrier protein synthase (AcpS). In order to obtain an activated FAS I in *E. coli*, we transformed *E. coli* with tagged *Mtb fas1* and *acpS* genes encoded by a separate plasmid.

We induced the expression of *Mtb* FAS I following induction of AcpS expression. FAS I was purified by Strep-Tactin affinity chromatography. Activation of *Mtb* FAS I was confirmed by the identification of a bound P-pant group on serine at position 1808 by mass spectrometry. The purified FAS I displayed biochemical activity shown by spectrophotometric analysis of NADPH oxidation and by CoA production, using the Ellman reaction. The purified *Mtb* FAS I forms a hexameric complex shown by negative staining and cryo-EM. Purified hexameric and active *Mtb* FAS I is required for binding and drug inhibition studies and for structurefunction analysis of this enzyme. This relatively simple and short procedure for *Mtb* FAS I production should facilitate studies of this enzyme.

## Introduction

*M. tuberculosis (Mtb)* possesses a unique lipid-rich cell wall that confers virulence, persistence and drug resistance. The important components of *Mtb* cell wall are α-alkyl-β-hydroxyl C80-90 fatty acids termed mycolic acids, and several polyketides (Pk) [1-4]. Several steps in mycolic acid biosynthesis were identified as drug targets for first-line anti-tuberculosis agents [5]. Mycobacteria unlike other prokaryotes or eukaryotes possess two types of fatty acid synthase systems, FAS I and FAS II. FAS I elongates acetyl-CoA to C16:0-26:0 fatty acids, the substrate for all subsequent steps of elongation and modifications.

FAS I is a multi-functional, multi-domain protein that is organized as a large (1.9 MDa) homohexameric complex [6,7]. Several structures of fungal FAS were reported, showing that the homologous fungal FAS is a α6β6 heterododecamer complex of 2.6 MDa which forms a barrel-shaped assembly with D3 symmetry. It has two reaction chambers, each containing three full sets of active sites that catalyze the entire acyl elongation reaction cycle, and are confined by a central wheel and two domes [8-10]. The mycobacterial FAS I is organized as a head to tail fusion of theα and β units of the fungal FAS [7]. The structure of *Mycobacterium smegmatis* (*M. smegmatis*) FAS was shown to resemble a minimized version of the fungal FAS with much larger openings in the reaction chambers. These architectural features of the mycobacterial FAS may be important for mycolic acid processing and condensing enzymes that further modify the precursors produced by FAS I[7].

The acyl intermediates produced during the fatty acid elongation process are attached covalently to an acyl carrier protein (ACP) domain of FAS I [9,11].

The ACP domain is positioned between the ketoreductase (KR) domain and the malonyl-palmitoyl transferase (MPT) domain and is activated through the transfer of a P-pant group from CoA to an ACP catalyzed by a 4’-PP transferase (PPTase) enzyme designated AcpS [7,11]. The *Mtb acpS* gene (*Rv2523c*) is located downstream of *fas* (*Rv2524c)*. It was shown that the *fas* and *acpS* genes are part of the same transcription unit forming a *fas1-acpS* operon under the control of FasR, an essential transcription regulator [12]. The functional association of AcpS of *Mtb* with FAS I is further supported by the structure-function organization of the analogous yeast FAS I, which possesses a PPTase domain - as part of the α unit of its α/β dodecamer complex [10,13] and which activates the ACP domain of the α subunit of the yeast FAS I complex [9,10,13]. However, it is not clear how the fungal PPTase domain on the α unit accesses the ACP domain which is far apart. It was suggested that the larger openings of mycobacterial FAS I allow easier access of AcpS to the ACP domain[7].

*Mtb fas1* was shown to be an essential gene and FAS I function was shown to be inhibited by pyrazinamide (PZA) analogs and pyrazinoic acid [14,15]. Pyrazinamide is a first line antituberculous drug and an indispensable component for the shortest curative regimen [16]. Thus, *Mtb* FAS I is a promising drug target against tuberculosis. Development of systems to obtain a recombinant active *Mtb* FAS I is a prerequisite for assays of drug binding and activity and for structure-function analysis.

So far, production of active *Mtb* FAS I was accomplished using the non-pathogenic strain *M. smegmatis* [17]. However, *fas1 g*ene of this strain was not fused to any tag, and thus sufficient purity of the protein required for structural studies has been challenging.

Herein, we show that sequential co-expression of *Mtb* AcpS and FAS I in *E.coli* followed by gentle purification of FAS I, results in binding of P-pant to a conserved serine residue of *Mtb* FAS I yielding an active hexameric FAS I complex.

## Materials and Methods

### Cloning of a recombinant *Mtb fas1* into pET100 D-Topo

A recombinant His tagged *Mtb fas1* was amplified from the plasmid pYUB955 (pMV206 plasmid bearing *Mtb fas1*) [14] using the primers HKNF and XbaR (Table 1) with Phusion^NEB^

**Table 1:**
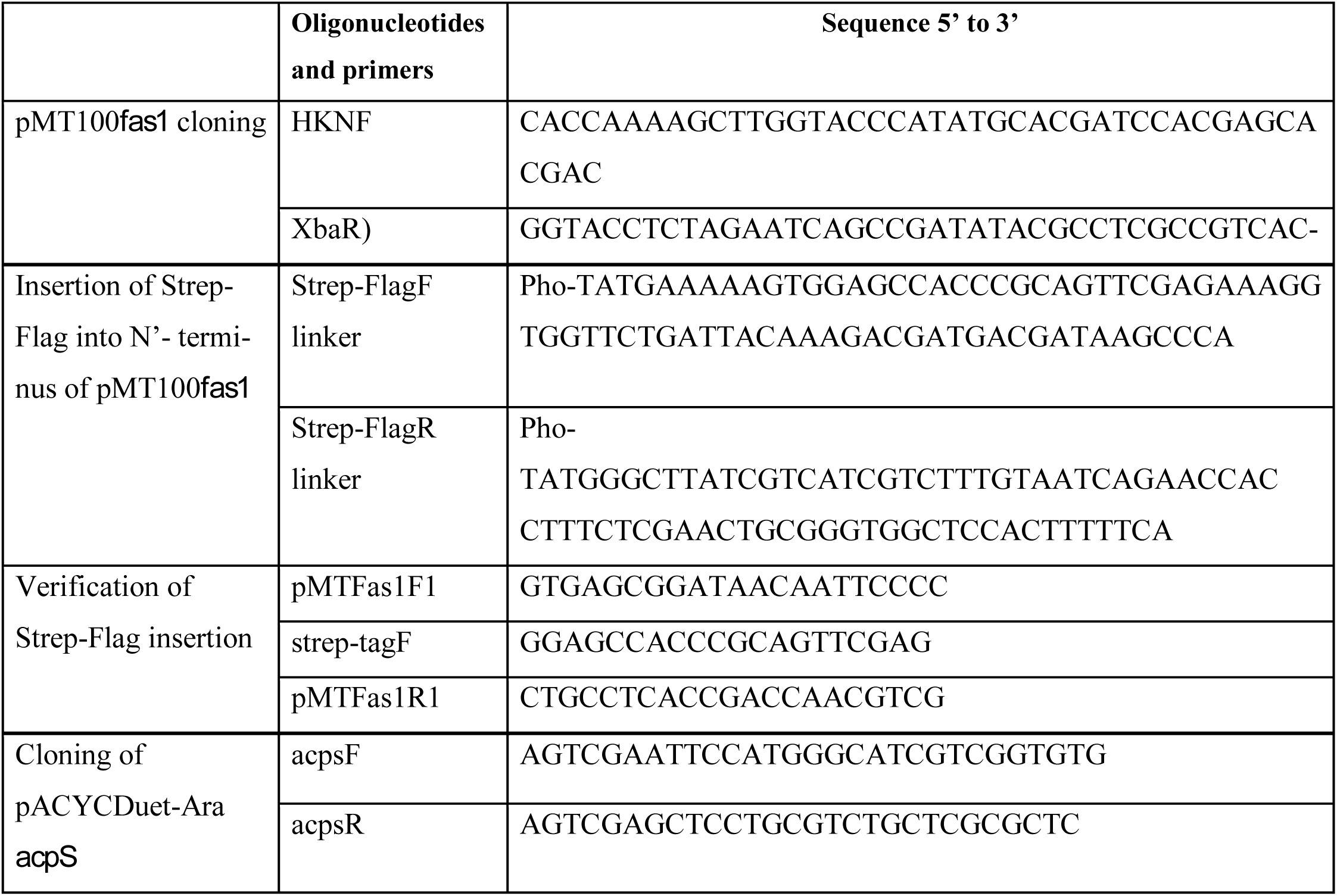
Oligonucleotides and primers for *Mtb fas1* and *acpS* cloning and modification.

### used in this study

The reaction was conducted at 99°C for 30 s and then run for 35 two-step cycles in a temperature of 99°C for 15 s and annealing and extension of 72°C for 5.5 min with final extension at 72°C for 10 min. The 10 kb PCR products were cloned into pET100 D-TOPO vector (Invitrogen-Thermo Fisher Scientific). Following transformation into DH5α *E. coli*, ampicillin resistant colonies were grown and the plasmids were purified and sequenced verifying the presence of the full sequence of *Mtb fas1* ORF. The resultant plasmid, designated pMT100-*fas1*, was used for all further steps described here.

### Construction of the *Strep-Flag-fas1* fusion plasmid

A Strep tag followed by Flag tag cassette was inserted onto the N-terminus of *Mtb fas1* in the expression vector pMT100-*fas1.* The DNA linker, harboring the cassette, was generated by annealing the two complementary primers, Strep-FlagF and Strep-FlagR, both phosphorylated at the 5’-end (Table 1). Annealing was performed by incubation of 1µM of each primer in a 0.2 ml tube placed into a PCR thermo block adjusted to 95^°^C for 5 min. Following the heating stage, the temperature was gradually reduced to ambient temperature. The double-stranded linker generated, harboring sticky ends at both ends for cloning into an NdeI site. The linker was then ligated into pMT100-*fas1* vector digested with NdeI and dephosphorylated. Correct orientation and integrity of the *Strep-Flag* insertion into pMT100*-fas1* were confirmed by PCR (Table 1) and DNA sequencing. The resulting vector was designated pMT100-*Strep-Flag*-*fas1*.

### Construction of AcpS expression vector controlled by arabinose induction

pACYCDuet-1 derivative vector harboring the arabinose control elements was constructed first. The arabinose control elements (*araC* repressor and BAD promoter) were amplified from the expression vector pBAD/gIII A (Invitrogen-Thermo Fisher Scientific) and inserted into the NarI and NcoI sites of the expression vector pACYCDuet-1, thus removing most of *lacI* gene and the T7 promoter from the first multiple cloning site. The resulting vector was designated pACYCAra. The *acpS* gene was amplified by using acpsF and acpsR primers (Table 1). pACYCAra vector and the PCR product harboring *acpS* were each digested with EcoRI and SacI followed by a ligation step. The integrity of pACYCDuet-Ara-*acpS* was confirmed by DNA sequencing.

### Expression and purification of *M. tuberculosis* FAS I

*Mtb* FAS I was co-expressed with *Mtb* AcpS to obtain activated FAS I protein. The plasmids pMT100-*Strep-Flag-fas1* and pACYCDuet-*Ara*-*acpS* were transformed into *E. coli* BL21 (DE3) cells, and selected on LB plates by resistance to ampicillin (AMP) and chloramphenicol (CAP) at 100 µg/ml and 34 µg/ml, respectively. An Amp^R^ CAP^R^ resistant colony was grown overnight in LB broth with these antibiotics. This culture was used to inoculate TYG broth containing 100 µg/ml ampicillin and 17 µg/ml chloramphenicol until an OD_600_ = 0.3-0.4 at 37°C was reached. Expression of AcpS was induced by addition of 0.2% arabinose and the culture was grown to OD_600_ = 0.6-0.8 at 37°C. The expression of FAS I was induced by addition of 0.5 mM IPTG at 15°C for 20 hrs. The cells were collected by centrifugation and resuspended in buffer A (100mM potassium phosphate buffer (KPB) pH 7.2, 150 mM KCl, 1 mM TCEP, 1 mM EDTA) containing protease inhibitor (cOmplete, Roche) 1 mM PMSF and DNase I (Roche) and lysed with a French press at 900 PSI. The lysate was centrifuged for 1 hr at 150,000 *g*. FAS I was precipitated from the lysate in two steps with 20% and 50% ammonium sulfate by overnight incubation and centrifuged for 20 min at 150,000 *g*. Precipitated FAS I was dissolved in buffer A and was loaded onto a 10 ml Strep-Tactin Sepharose beads (GE Healthcare) gravity flow affinity column (BioRad) [18]. The column was washed with buffer A, and eluted with buffer A containing 2.5 mM desthiobiotin. FAS I containing fractions were collected and concentrated using 15ml 50K Amicon^®^ filter units with a MW cut off of 50 kDa, and flash frozen by liquid nitrogen and stored at −80°C. For some samples, we analyzed this purified FAS I by size exclusion chromatography using a Superose 6 10/300GL (GE Healthcare) equilibrated with buffer A.

Purification of *Mtb* FAS I from *M. smegmatis* Δ*fas1* (*attB::Mtb fas1*) strain, designated mc^2^ 2700 was used for comparison and is described in the supporting information (S method).

### Sample preparation for mass spectrometry

The following purified recombinant FAS I samples were examined: FAS I from *M. smegmatis* mc^2^ 2700 purified as described in S method; and the fusion protein Strep-Flag-FAS I expressed in *E.coli* BL21 DE3 cells with or without expression of AcpS. The proteins were loaded onto an 8% SDS PAGE gel, each showing the band of 326 kDa FAS I monomer. The stained bands associated with FAS I, were sliced and placed in 0.65 ml silicon tubes (PGC Scientific), and destained with 25 mM NH_4_HCO_3_ in 50% acetonitrile (ACN) and completely dried. The samples were reduced in with 10 mM DTT in 25 mM NH_4_HCO_3_ at 56°C for 1 hr and alkylated with 55 mM iodoacetamide in 25 mM NH_4_HCO_3_ in the dark for 45 min at room temperature (RT). The dried gel slices were then washed with 25 mM NH_4_HCO_3_ in 50% ACN and dehydrated, followed by trypsin digestion with 12.5 ng/μL trypsin in 25 mM NH_4_HCO_3_ at 4°C for 10 min, and followed by overnight incubation at 37°C. The reaction was quenched by adding 1:1 volume of trifluoroacetic acid (1%). The eluent was acidified with formic acid, dried and kept at −80°C for analysis.

### Identification of the P-pant group by Mass Spectrometry

Gel samples were analyzed using an Orbitrap Fusion Lumos coupled to a nanoAcquity naboUPLC (Waters, Milford, MA, USA). Samples were trapped on a Pepmap C18 trap column (100 μm internal diameter, 20 mm length, 5 μm particle size; Thermo Fisher Scientific), and separated on an Easy-Spray Pepmap RSLC column (75 μm internal diameter, 250 mm length, 2μm particle size; Thermo Fisher Scientific) at a flow rate of 0.35 μL/min, using a gradient of 4-20% solvent B (80% ACN) in 50 min, followed by 5 min in 20-30% solvent B.

Data were acquired in Data Dependent mode, using a 3 sec Top speed method. MS1 resolution was set to 120,000 in the orbitrap (at 200 m/z), maximum injection time was set to 50 msec automatic gain control (AGC) target was 4e^5^. High priority MS2 scans were set to Collision Induced Dissociation (CID) in the Orbitrap (35 normalized collision energy (NCE) 10 msec, activation Q 0.25) at 15000 resolution (at 200m/z). Lower priority scans were performed with electron-transfer/higher-energy collision dissociation (EThcD) in the Orbitrap (60 msec reaction time of electron-transfer dissociation (ETD) with 3e^5^ AGC reagent target, 18NCE) at 15000 resolution (at 200m/z), with preference of high charge ions. In case that the pantethinylacetamide ejection ion (318.1488 m/z) was detected in the CID scans two additional fragmentation events were triggered. The first event triggered higher energy collision dissociation (HCD) fragmentation of the ejection ion itself to verify its identity as pantethinylacetamide based on indicatory ions as described [19]. MS3 was performed with 30NCE HCD in the Ion Trap at normal speed, with first mass set to 85 m/z (AGC target of 3e^4^, 256 msec fill time). Second, the same precursor was sent to another MS2 scan, EThcD scan, in the orbitrap – ETD reaction was set to 60 msec with 3e^5^ AGC target, followed by 20NCE HCD, scanned at 15000 resolution (at 200 m/z). Scan range was set to 120-1600 m/z, allowing 100 msec accumulation time.

Data analysis was performed using Byonic search engine (Protein Metrics Inc), against a concatenated database of *E. coli* and *M. tuberculosis*, setting carbamidomethylation on Cysteine allowing oxidation on Methionine, deamidation on Glutamine and Aspargine and phophopantetheinylacetamide on Serine, searching only the EThCD spectra. Identification and the associated MS3 scans were verified manually.

### FAS I activity measurements

FAS I activity was monitored as a decrease in absorbance at 340 nm at RT for 20 min resulting from NADPH oxidation in each C2 elongation during fatty acid synthesis [17]. The assay was conducted in 0.1 ml of KPB (pH 7.5, 0.2M), using NADPH at 200 µM, acetyl-CoA at 400 µM, and malonyl-CoA at 200 µM concentration. Twenty five to fifty micrograms of purified FAS I were added to the enzymatic assay. We also followed specific inhibition by 5-ClPZA, mycobacterial FAS I inhibitor [14], at a concentration of 200 µg/ml, to validate the specificity of the observed NADPH oxidation to *Mtb* FAS I activity. In addition, FAS I activity was monitored using free CoA accumulation under the same reaction conditions with 5,5’-dithiobis-(2-nitrobenzoic acid) (DTNB, Ellman’s reagent) [20] at 20 µM, by changes in the absorbance at 412 nm for 20 min. One unit of enzyme activity was defined as the amount of enzyme required to catalyze the oxidation of 1 µmol of NADPH per minute. Specific activity based on NADPH oxidation was expressed in nmole/min i.e. as milli-units (mU). Specific activity was calculated as the difference in OD at 340 nm in 20 minutes divided by the extinction coefficient for NADPH, 6220 molar/cm and then divided by mg of protein used in the reaction and by 10^4^, to account for the reaction volume of 0.1 ml.

### EM studies

For negative stain EM, 3.5 μL of purified FAS I sample at 0.4 mg/ml concentration was applied to glow-discharged, homemade 300 mesh carbon-coated copper TEM grids for 30 sec. The grids were stained with 2% uranyl acetate. Samples were visualized in an FEI Tecnai T12 TEM operated at 120 kV, equipped with a Gatan OneView camera. For cryo EM, 3.5 μL of purified FAS I solution at 0.4 mg/ml concentration was applied to glow-discharged Quantifoil holey carbon grids (R2/2, 300 mesh) coated with a thin layer of carbon. Grids were plunged into liquid ethane cooled by liquid nitrogen, using a Leica EM-GP plunger (3.5 sec blotting time, 80% humidity). Grids were imaged at liquid nitrogen temperature on an FEI Tecnai TF20 electron microscope operated at 200 kV with a Gatan side entry 626 cryo-holder. Images were recorded on a K2 Summit direct detector (Gatan) mounted at the end of a GIF Quantum energy filter (Gatan). Images were collected in counting mode, at a calibrated magnification of 23,657 yielding a pixel size of 2.11 Å. The dose rate was set to ∼8 electrons per physical pixel per second and a total exposure time of 10 sec, resulting in an accumulated dose of ∼18 electrons per Å^2^. Each image was fractionated into 40 sub-frames of 0.25 sec. Defocused range was 1.2 to 4 μm. All dose-fractionated images were recorded using an automated low dose procedure implemented in SerialEM10 [21]. Recorded image frames were subjected to whole image beam-induced motion correction using MOTIONCORR [22]. Contrast transfer function parameters were estimated using CTFFIND3 [23]. Twenty nine images were used, from which 1906 particles were manually picked using EMAN2 e2boxer [24]. Particles were extracted into 180×180 pixel boxes and 2D classified into 6 classes with a round mask of 300 Å diameter using RELION 1.4 [25].

## Results

### Purification and characterization of recombinant active *Mtb* FAS I

The recombinant *Mtb* FAS I which was expressed following *Mtb* AcpS expression in *E. coli* Bl21, yielded about 0.4 mg of pure recombinant *Mtb* FAS I per one liter of TYG broth. The yield for *Mtb* FAS I purification from *M. smegmatis* mc^2^ 2700 (S method) was about 0.15 mg/liter. The procedure from cell lysis to final, purified FAS I protein was conducted in two days and in 4-5 days, for the recombinant FAS I in *E. coli* and *M. smegmatis* mc^2^ 2700, respectively. Following purification by Strep-Tactin affinity column, purified FAS I migrated on SDS PAGE as 326 kDa, corresponding to a FAS I monomer (Fig 1A). A band from SDS PAGE gel was submitted to analysis by mass spectrometry following tryptic digest that confirmed its identity as *Mtb* FAS I (S1 Fig)

**Fig 1:**
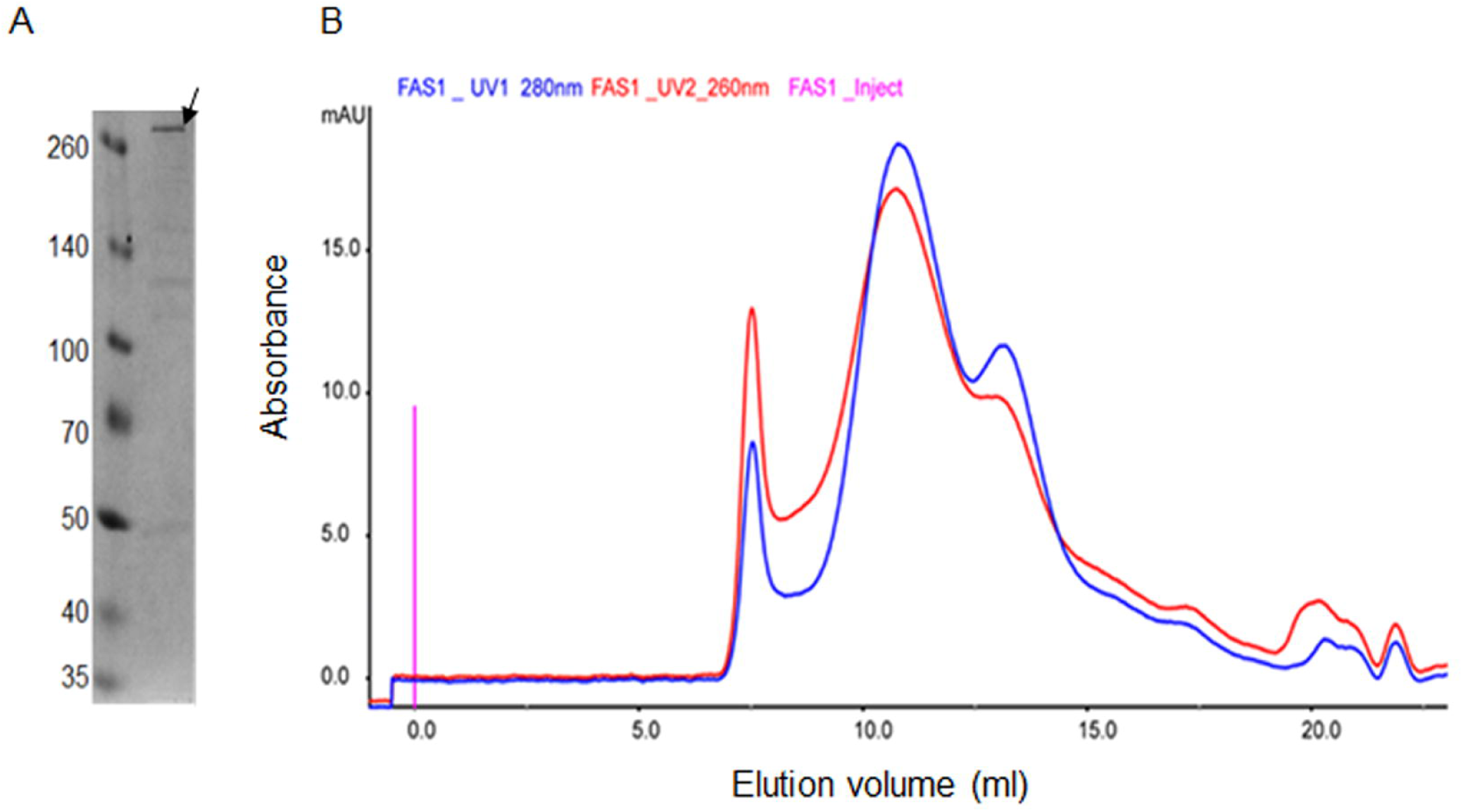
Analysis of purified active recombinant *Mtb* FAS I following elution from Strep-Tactin affinity column: A. SDS –PAGE (8%) gel analysis, FAS I is marked by arrow. B. Migration profile chromatogram of *Mtb* FAS I following analysis on superose 6 column.

This migration profile through superpose 6 column (Fig 1B) is in agreement with an assembled large hexameric complex as examined by EM. In some preparations, the size exclusion profile using superpose 6 column, exhibited two peaks. One peak migrated at 11.47 ml and a second peak migrated as a smaller complex at 12.7 ml. Both peaks correspond to pure FAS I as shown by SDS PAGE but only the 11.47 ml peak exhibited hexameric structures in EM and enzymatic activity. It is likely that the protein migrating at 12.7 ml are incompletely assembled FAS I complexes.

### Confirmation of P-pant binding to FAS I

To identify the P-pant group binding to serine as a validation of FAS I activation by AcpS, we conducted a bottom proteomic analysis of a tryptic digested gel band. Identification of the modified peptide was performed using a specialized search engine and was confirmed by two distinct measures: the first is MS3 of the supposedly pantetheinylacetamide ejection ion, producing indicative ions of its fragments, which was inspected manually based on the preceding findings of Meier JL et al [19]. The second is manual inspection of the fragmentation spectra of the peptide itself. Expression of FAS I without *Mtb* AcpS did not yield any peptides bearing P-pant group on Serine 1808 (Fig 2A). We identified a single P-pant modification site in the activated FAS I expressed in *E. coli*, on serine 1808 (Fig 2B), as was observed for *Mtb* FAS I expressed in *M. smegmatis*. (S3 Fig.)

**Fig 2:**
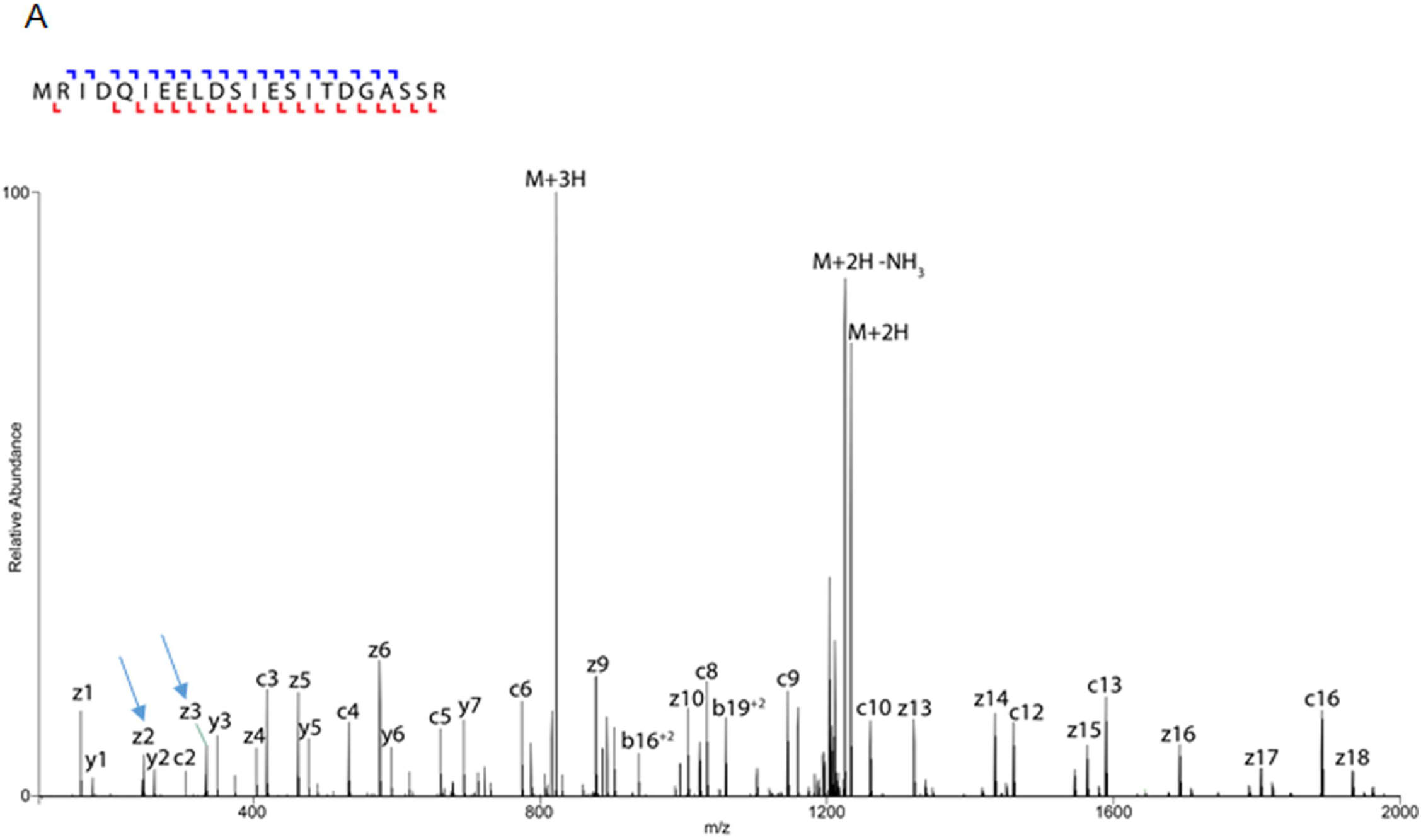
Annotated spectra of P-pant carrying peptide from StrepFlag *Mtb* FAS I expressed in *E. coli* Bl21 without AcpS and following AcpS expression.

An electron-transfer/higher-energy collision dissociation (EThcD) spectra acquired for a P-pant carrying peptide from FAS I protein. Identified ions are annotated on the spectrum, as well as on the sequence presented on top of the figure. P-pant modification is accurately localized at serine 1808 as show by the identification of the flanking backbone fragments. (indicated by arrow at z3 vs. z2). A. peptide spectrum without AcpS expression B. peptide spectrum following AcpS expression.

### FAS I activity

We utilized spectrophotometric assay of NADPH oxidation resulting in decrease in absorbance at 340 nm. This is a commonly used indirect assay for the study of fatty acid synthesis and elongation. A decrease in 340-nm absorbance was shown only for the AcpS-activated FAS I (Fig. 3A) bearing P-pant.

**Fig 3:**
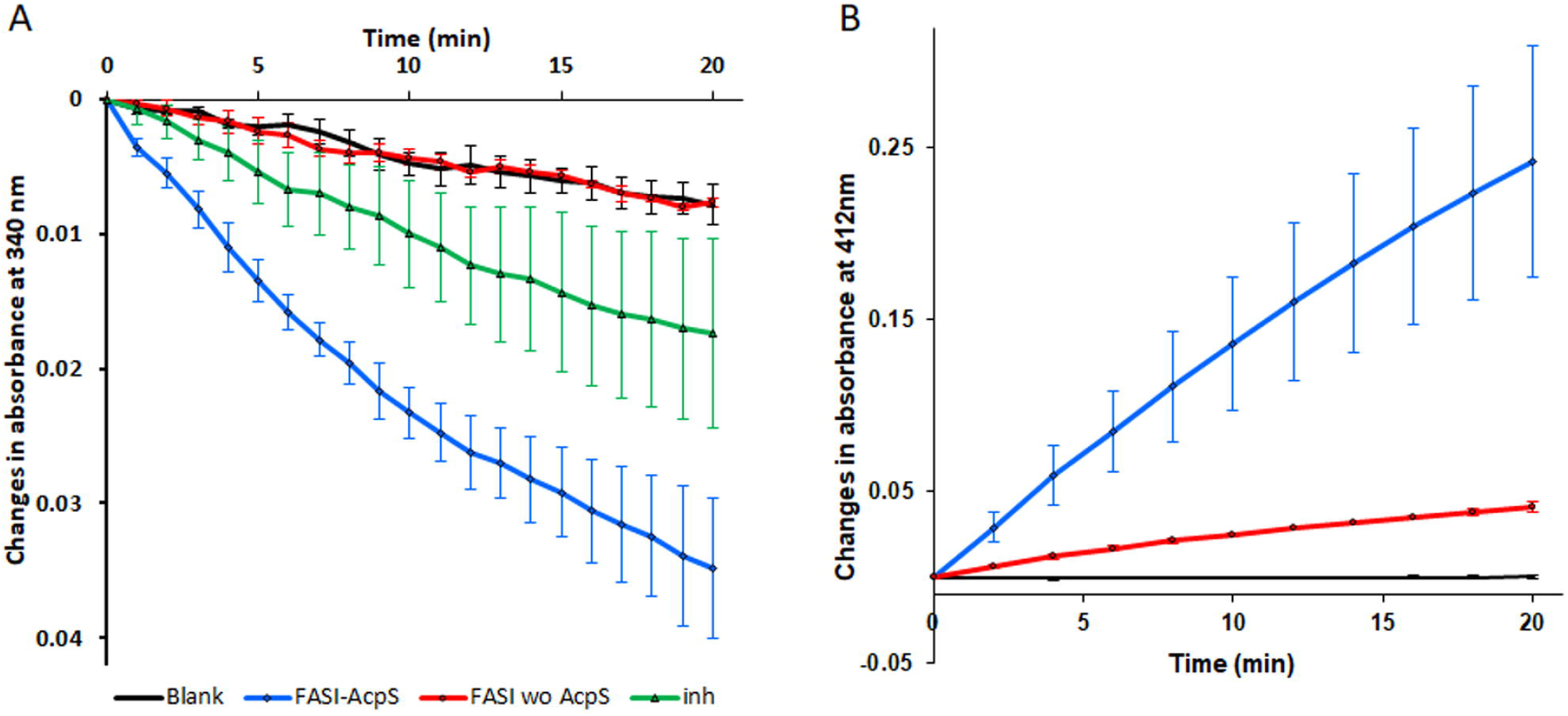
Recombinant *Mtb* FAS I activity.

A. NADPH oxidation assay: blank denotes no enzyme, FAS I denotes FAS I without AcpS, FAS I-AcpS denotes FAS I following AcpS expression, inh denotes FAS I following AcpS expression with 5-ClPZA at 200 µg/ml. B. Ellman’s reaction: blank, FAS I, and FAS I-AcpS as in A. Each plot represent two triplicate batch experiments. Error bars represent standard errors.

The release of free CoA (yielding free thiol groups), as predicted from malonyl CoA and acetyl CoA binding to ACP during fatty acid synthesis and elongation, was observed using the Ellman’s assay resulting in increased absorbance at 412 nm (Fig. 3B). FAS activity in both assays was dependent on Acetyl CoA and was inhibited by addition of 5-Cl-PZA, a specific inhibitor of FAS I (Fig 3A).

FAS I activity was observed only for the hexameric fraction of FAS I following superpose 6 elution (data not shown). We quantified the specific activity as the consumption of NADPH with two different batches of purified *Mtb* FAS I, each batch in triplicate, yielding activity of 1.75 ± 0. 18 and 1.28 ± 0.195 mU /mg respectively. This activity was lower than what was previously measured with a recombinant *Mtb* FAS I isolated from *M. smegmatis* [17]. The specific activity of FAS I purified herein from *M. smegmatis* mc^2^ 2700 (S method and S2 Fig). was 4.88± 0.21 mU/mg

### EM studies

Purified active FAS I from *E.coli* was analyzed by transmission electron microscopy (TEM) in order to assess the assembly state of the complex. The purified sample was first observed by negative stain EM (Fig 4A).

**Fig 4.**
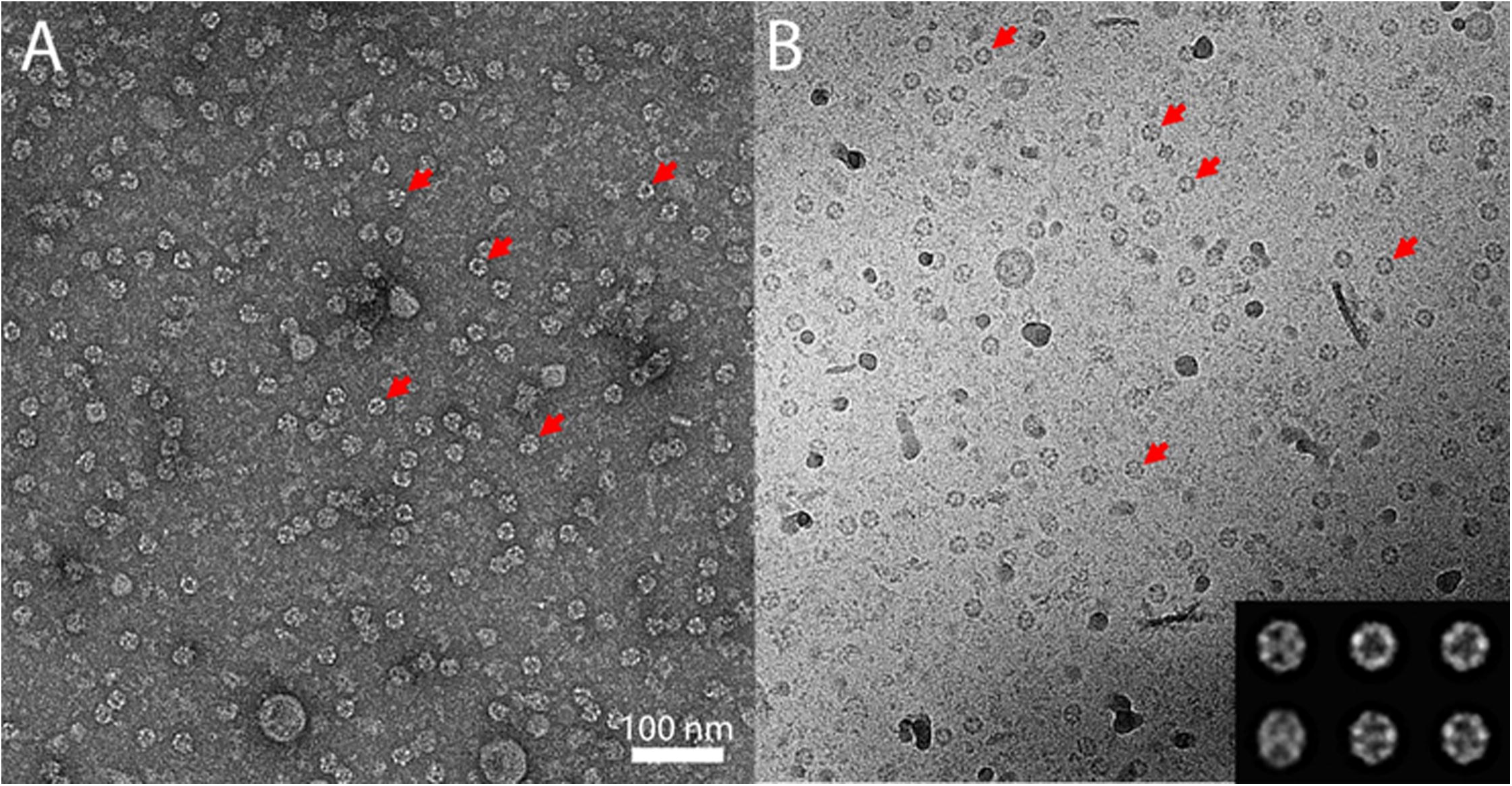
Negative stain (A) and cryo TEM (B) images of FAS I. Sample showing assembled complexes, red arrows point to some of the assembled FAS I complexes. Class averages of the cryo-EM FAS I complexes are shown in the inset.

TEM images clearly show assembled complexes at the expected size which dominate the grid. FAS I assembles into a barrel-shaped complex with D3 symmetry. The observed rings with a diameter of 22 nm in the negative-stain images correspond to top views of the complex, along the 3-fold symmetry axis, and elliptical-shaped complexes correspond to tilted views. The same ring and elliptical-shaped complexes were also observed under cryogenic conditions (Fig 4B). Plunge-frozen samples at liquid nitrogen temperatures are less prone to artifacts because they are hydrated, and the contrast results from the organic material rather than from heavy metal stain. A data set of 1906 FAS I complexes was manually picked from cryo-TEM images and classified in 2D. Class averages clearly show 3-fold symmetry in the top views, as well as the expected tilted views (Fig 4B inset). We note that the data set lacks side views, likely because of preferential absorbance of the cavity opening to the carbon.

## Discussion

This study describes the expression of a 4’-phosphopantetheinylated active *M. tuberculosis* fatty acid synthase I in *E. coli* following sequential expression of *Mtb* FAS I after its *Mtb* PPTase enzyme, AcpS.

Structure-function studies of mycobacterial FAS I require pure and homogeneous preparations of properly assembled complexes. It is possible to purify native FAS I from mycobacteria species but such purification of activated FAS I is a laborious procedure, requiring a combination of ammonium sulphate precipitation, anion exchange and size exclusion chromatography or sucrose cushion and gradient separations [7,17]. In addition, cultivating species of mycobacteria, especially large volumes is more demanding and a lengthy procedure due to the slower growth of mycobacteria.

Alternatively utilizing *E. coli* for producing activated mycobacterial FAS I, and the subsequent rapid purification procedure based on Strep -Tactin affinity capture, effectively mitigates the above mentioned limitations. Although the yield of this FAS I purification is modest, upscaling is feasible.

The activity of *Mtb* FAS I was shown to be dependent on activation by AcpS resulting in binding of P-pant group to Ser1808, a highly conserved serine in FAS I of other mycobacteria and corynebacteria species [26]. Proteomic detection of P-pant modified peptides is difficult due to their low abundance and the labile nature of the modification which leads to a partial loss of the side chain. This in turn, impairs the scoring of possible identification by search engine. Thus, the identification of the P-pant modification was improved by combining traditional search engines with further validation of P-pant via identification of the P-pant ejection ion by using MS3, in addition to ETD fragmentation which preserves the intact modification [19]. P-pant identification in this study was finally validated by manual inspection of the fragmentation spectra of the peptide itself as described above, but without enrichment steps of the purified FAS I. FAS I expression without AcpS, yielded inactive FAS I that does not have P-pant on serine at position 1808, suggesting that native AcpS from *E. coli* is incapable to activate *Mtb* FAS I. However, the specific activity of the recombinant FAS I purified from *E.coli* using the same procedure to measure FAS I activity, was lower than the activity of *Mtb* FAS I purified from *M. smegmatis* mc^2^ 2700 strain. We assume that the lower specific activity of the *E. coli* expressed FAS I compared to mycobacterium, can be attributed to two factors: incomplete activation of all FAS I molecules upon co-expression of *Mtb* FAS I with its AcpS and lower stability of the FAS I hexamer complex, following propagation in an *E. coli* host rather than in mycobacterium species. Nevertheless, only FAS I activated by AcpS exhibited specific activity which was inhibited by 5-ClPZA.

In fungi, the pantothenic transferase (PPT) and its substrate, ACP domain, are both part of the α unit. The PPT is on at the C-terminus and the ACP domain is at the N-terminus, being separated from each other by the ketoreductase (KR) and ketoacyl synthase (KS) domains. Moreover, the structure of fungal FAS revealed a barrel-shaped architecture in which PPT is located at the outside wall and thus spatially separated from the ACP which is located at the inner space. This finding leads to the conclusion that activation of fungal ACP likely takes place prior to the assembly of the dodecameric 6α6β complex in a conformation that enables PPT to activate ACP [10,27]. In mycobacteria, AcpS encoded by the *acpS* gene is a part of the *fas1-acpS* operon, and is not included in the FAS I complex. In view of the possible limited access of AcpS to ACP within the reaction chamber, we chose to use sequential co-expression of AcpS and FAS I to enable activation of ACP prior to the formation of the homo-hexameric complex. A parallel co-expression of *Mtb* AcpS and FAS I rather than sequential coexpression as we applied, was described for FAS I (designated FAS B) from *Corynebacterium eficiens*. FAS I enzyme of this species displays 52% homology and 66% similarity (4% gaps) to *Mtb* FAS I and was reported to yield crystals that diffracted X-ray to about 4.5 Å resolution [28]. So far, we could not obtain crystals of the purified active *Mtb* FAS I herein. TEM images of the purified FAS I clearly show that the complexes are stably assembled displaying the expected size and shape as described. Although some protein particles are seen at the background, both of the negative stain and cryo-TEM images are dominated by the assembled complexes at the expected size. The dominance of top views is due to preferential absorbance of one particular surface of the particle to the carbon support. Specifically, the opening of the FAS I particle cavity has higher affinity to the carbon support compared to other surfaces of the particle.

This relatively simple and shorter procedure for *Mtb* FAS I production through sequential coexpression of *Mtb* AcpS and FAS I in *E.coli*, should facilitate studies of this essential drug target, enzyme complex.

## Supporting information

**S Method. Purification of *Mtb* FAS I from *M. smegmatis* mc^2^ 2700** Δ***fas1* (*attB::Mtb fas1*) strain**

**S1 Fig. Mass spectrometry analysis of the recombinant *Mtb* FAS I purified from *E. coli* Bl21 following tryptic digest.**

**S2 Fig**. ***Mtb* FAS I purified from *M. smegmatis***. A. Migration profile chromatogram of FAS I purification on Superose 6 column. FAS I migrates at 11.4ml (blue - absorbance at 280nm, red – absorbance at 260nm, pink – injection point). B. Coomassie blue stained 6%/15% SDS PAGE of 11.4ml peak from (A).

**S3 Fig. Annotated spectra of P-pant carrying peptide of *Mtb* FAS I purified from *M. smegmatis* strain mc^2^ 2700.** An electron-transfer/higher-energy collision dissociation (EThcD) spectra acquired for a p-pant carrying peptide from FAS I protein. Identified ions are annotated on the spectrum. P-pant modification is accurately localized at serine 1808 as shown by the identification of the flanking backbone fragments (indicated by arrow at z3 vs. z2)

## Acknowledgements

We are grateful to the continuous support of the Kimmelman Center for Macromolecular Structure and Assembly at the Weizmann Institute of Science.

